# The exosomes derived from urine of premature infants target MAPK8 via miR-30a-5p to protect cisplatin-induced acute kidney injury in mice

**DOI:** 10.1101/2021.08.29.458088

**Authors:** Mingming Ma, Qiao Luo, Lijing Fan, Wei-Long Li, Qiang Li, Yu Meng, Chen Yun, Hongwei Wu, Yongping Lu, Shuang Cui, Fanna Liu, Bo Hu, Baozhang Guan, Huanhuan Liu, Shengling Huang, Wenxue Liang, Stanislao Morgera, Bernhard Krämer, Shao-Dong Luan, Lianghong Yin, Berthold Hocher

## Abstract

**Aim:** Acute kidney injury (AKI), a global public health issue, not only causes millions of deaths every year, but is also a susceptible factor for chronic kidney disease (CKD). Nephrotoxic drugs are an important cause of AKI. There is still a lack of effective and satisfactory prevention method in clinical practice. This study investigated the protective effect of the exosomes derived from urine of premature infants on cisplatin-induced acute kidney injury.

**Methods:** Isolation of exosomes from fresh urine of premature infants: The characteristics of exosomes were determined by flow cytometry, transmission electron microscopy and Western blotting. A C57BL/6 mice model of cisplatin-induced acute kidney injury was established. The mice in the experimental group were given 100ug exosomes dissolved in 200ul solution. The mice in the control group were given normal saline (200ul). These treatments were performed 24 hours after AKI was induced by intraperitoneal injection of cisplatin. To evaluate renal function, blood was drawn 24 hours after AKI model was established and serum creatinine (sCr) was measured. The mice were euthanized 72 hours after exosome treatment. The kidneys were collected for pathological examination, RNA and protein extraction, and the evaluation of renal tubular damage and apoptosis. In the in-vitro experiment, human renal cortex/proximal tubular cells (HK2) was induced by cisplatin to assess the protective ability of the exosomes derived from urine of premature infants. Quantitative reverse transcription polymerase chain reaction (qRT-PCR) and Western Blotting were used to evaluate the effect of exosomes treatment on the apoptosis of HK2 cells induced by cisplatin. Exosome microRNA sequencing technology and bioinformatics analysis method were applied to investigate the miRNAs enriched in exosomes and their target genes. The dual luciferase gene reporter system was used to detect the interaction of target genes.

**Results:** Treatment of exosomes derived from urine of premature infants could decrease the level of serum creatinine and the apoptosis of renal tubular cell, inhibit the infiltration of inflammatory cell, protect mice from acute kidney injury induced by cisplatin and reduce mortality. In addition, miR-30a-5p was the most abundant miRNA in the exosomes derived from urine of premature infants. It protected HK2 cells from cisplatin-induced apoptosis by targeting and down-regulating the 3’UTR of mitogen-activated protein kinases (MAPK8) mRNA.

**Conclusions:** According to our results, the exosomes derived from urine of premature infants alleviated cisplatin-induced acute kidney injury in mice and inhibited the apoptosis of human proximal tubular cells (HK2) induced by cisplatin in vitro. MiR-30a-5p in exosomes inhibited cisplatin-induced MAPK activation, ameliorated apoptosis, and protected renal function. The exosomes derived from urine of premature infants provided a promising acellular therapy for AKI.

## Introduction

As a clinical syndrome resulted from various etiology and pathophysiological processes, acute kidney injury (AKI) is characterized by sudden decline (1-7d) and continuous deterioration (>24h) of glomerular filtration function [1, 2]. AKI is a common disease with an incidence rate of 2100/ million people worldwide. It is more commonly seen in hospitalized patients, with an estimated incidence of 10%-15% [2]. In the past, AKI was considered to be a self-limiting and completely reversible disease [1, 3]. In recent years, more and more studies have suggested that not all types of AKI were reversible[4]. Now, it has been acknowledged that the acute change in renal function was associated with long-term prognosis, including progression to chronic kidney disease and cardiovascular disease, persistent dysfunction, and even death[1]. As an episode leading to chronic kidney disease (CKD) or end stage renal disease (ESRD)[3], AKI has an increasing incidence year by year. Its impact on long-term human health is far greater than we have recognized. Although the prevention and early treatment of AKI are constantly improved, the prognosis of AKI is still poor. There is still a lack of effective therapeutic strategy[5, 6].

Among the new treatment methods of AKI, extracellular vesicles (EVs) derived from stem cells have attracted more and more attention. Accumulating evidence showed that exosomes derived from mesenchymal stem cells (MSC) carried and secreted the “label signals” of miRNA, which was a key factor influencing therapeutic effects [7–9]. Extracellular vesicles play a role in multiple regeneration processes, including inducing the survival and proliferation of renal tubular cell, and inhibiting apoptosis, inflammation, and fibrosis [10, 11]. After EV enters the target cell, the transfer of its content (such as protein, RNA and DNA) is the foundation of regulating the cell fate[12]. Previous studies have found that the miRNAs transferred by MSC-EVs was considered to be the main effector promoting the proliferation and survival of tubular cell[13]. Interestingly, human urine-derived stem cells (hUSCs) isolated from human urine were reported to secrete proangiogenic growth factors to maintain the differentiation potential of endothelial cells, which played a pivotal role in the treatment of diabetes[14]. A previous study showed that the exosomes secreted by human urine-derived stem cells could improve diabetic kidney injury by inhibiting apoptosis of podocyte and promoting angiogenesis[15], Yu-Rui Duan et al. further found that miR-16-5p secreted by urine-derived exosomes could inhibit the damage to podocyte induced by high glucose, and thus ameliorating diabetic kidney injury [16]. In addition, recent studies reported that miR146a-5p and Klotho in EVs derived from healthy adult urine attenuated renal tubular cell apoptosis and promoted cellular repair in acute kidney injury model[17, 18]. Therefore, urine-derived exosomes may be a promising method for kidney repair, but the specific mechanism is not yet fully understood. This study aimed to investigate the protective effect of the exosomes derived from urine of premature infants on cisplatin-induced acute kidney injury. In addition, we explored the possible mechanisms of this protective effect by analyzing the miRNAs in the exosomes derived from urine of premature infants.

## Methods and Materials

### Isolation and identification of the exosomes derived from urine of premature infants

This study was approved by the Ethics Committee of Jinan University (ethics application number 2020724-01, approval number IACUC-20200813-05). The informed consent was obtained from the guardian of the urine donor. We used fresh and sterile mid-section urine samples of healthy full-term infants (37 weeks ≤ gestational age <42 weeks) and premature infants (gestational age ≤ 36 weeks) in the newborn ward of our hospital. Each urine sample was about 15ml. The sterile urine collection bag was placed at the external urethral orifice to collect urine within 1 day after birth. Fecal contamination should be carefully avoided during the collection, and then the urine sample was transferred to the 20ml sterile test tube. Exosomes were isolated as previously described[17, 19]. After the sample was collected, the urine sample was immediately centrifuged at 2,000×g in 4°C for 10 minutes by using the German Eppendorf centrifuge to remove cells and debris. The supernatant was transferred to a new test tube, ultracentrifuges at 17000 × g in 4°C for 45 minutes, and then filtered through a 0.22um sterile filter. and take the supernatant. The supernatant was centrifuged by using a Beckman Coulter ultracentrifuge at 200,000 g in 4°C for 70 minutes. The supernatant was removed. The remaining pellet was suspended in phosphate buffer saline (PBS), and then centrifuged at 200,000 g in 4°C for 70 minutes. The supernatant was discarded, and the exosome particles from each sample were suspended in 50μL RNase-Free water and stored in −80°C refrigerator for further use. All methods were carried out in accordance with relevant laboratory guidelines and institutional regulations. The protein concentration of exosomes was determined by using the BCA protein determination kit (BIO-RAD, 500-0201). Western blotting and flow cytometry were used to detect exosome markers. The morphology of exosomes was observed under the transmission electron microscope (TEM), and the average diameter of exosomes was measured. The particle size and concentration were measured with ZetaView PMX110 (Particle Metrix, Meerbusch, Germany), and ZetaView 8.04.02 was used to analyze the data.

### Isolation and high-throughput sequencing of miRNA

SeraMir Exosome RNA Purification Column kit (SBI) was used for extracting total RNA in exosomes. The quality and purity of the extracted RNA were detected by using Agilent 2100 bioanalyzer. Ion Total RNA-Seq kit V2 (Thermo Fisher) was used to construct a small RNA sequencing library. Sequencing was conducted on the Ion Proton sequencing platform. The ACGT101-miRv4.2 (LC Sciences) software program was used to analyze high-throughput sequencing data. DEGseq (R software language package) and perl script was used to perform statistical analysis on the identified miRNAs.

### Western blot

Tris-glycine gel (10%) was used to separate 30ug protein in urinary exosomes from premature infants. The protein was then transferred onto the polyvinylidene fluoride (PVDF) membrane. Protein gel electrophoresis and transfer were performed in accordance with standard protocols. Skim milk (5%) was used to block the membranes at room temperature for 1 hour to remove non-specific binding. The membranes were incubated with the diluted primary antibodies (CD63 1:2000, CD81 1:1000, CD9 1:1000) overnight at 4°C. The membranes were rinsed 3 times with TBST buffer. Secondary antibody (IgG 1:2000) was then incubated with the membranes at room temperature for 2 hours followed by rinsing with TBST buffer. SuperSignal West Femto chemiluminescence reagent was added onto the PVDF membrane. The chemiluminescence imaging analyzer was applied to observe and take photographs of the membranes. The antibodies employed in the experiment are listed in Table 1.

**Table 1.**
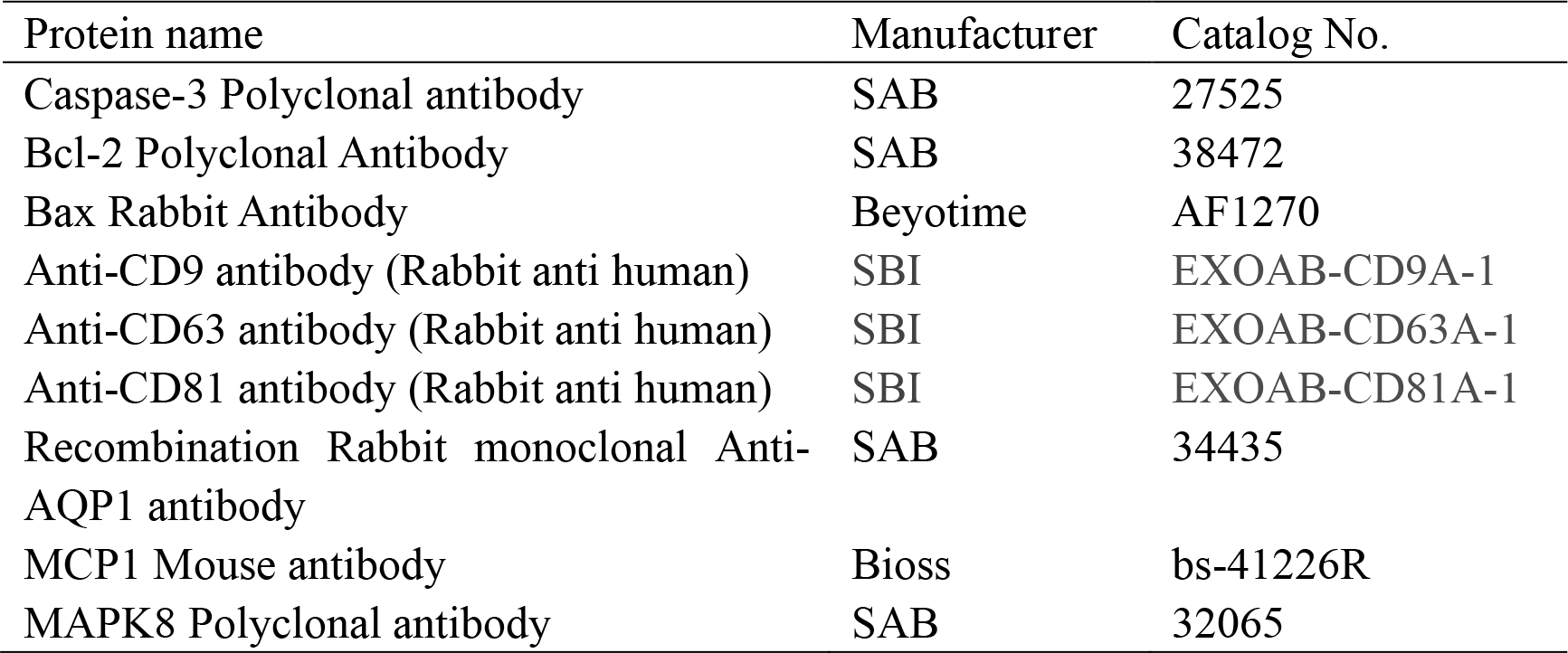
Antibodies used in this study

| Protein name | Manufacturer | Catalog No. |
| --- | --- | --- |
| Caspase-3 Polyclonal antibody | SAB | 27525 |
| Bcl-2 Polyclonal Antibody | SAB | 38472 |
| Bax Rabbit Antibody | Beyotime | AF1270 |
| Anti-CD9 antibody (Rabbit anti human) | SBI | EXOAB-CD9A-1 |
| Anti-CD63 antibody (Rabbit anti human) | SBI | EXOAB-CD63A-1 |
| Anti-CD81 antibody (Rabbit anti human) | SBI | EXOAB-CD81A-1 |
| Recombination Rabbit monoclonal Anti-AQP1 antibody | SAB | 34435 |
| MCP1 Mouse antibody | Bioss | bs-41226R |
| MAPK8 Polyclonal antibody | SAB | 32065 |

### qRT-PCR analysis

qRT-PCR was performed by using 2XTaqMan qPCR Master Mix (solarbio) according to the instructions of the kit. The reaction mixture contained 1.33ul cDNA solution, 10ul mixed solution, 7.67ul RNA enzyme-free water, and 1ul forward or reverse primer. QRT-PCR was carried out on the CFX96 touch instrument. the cycle scheme was set as follows: PCR reaction program: 95℃,10 minutes, warm boot; 95℃,15 seconds; 60℃, 1 minute, amplification for 40 cycles; deionized water was set as negative contrast. Ct values were calculated by using the thresholds and baselines set automatically. Ct values higher than 38 were excluded from the analysis. The primers used in qRT-PCR were listed in Table 2.

**Table 2.**
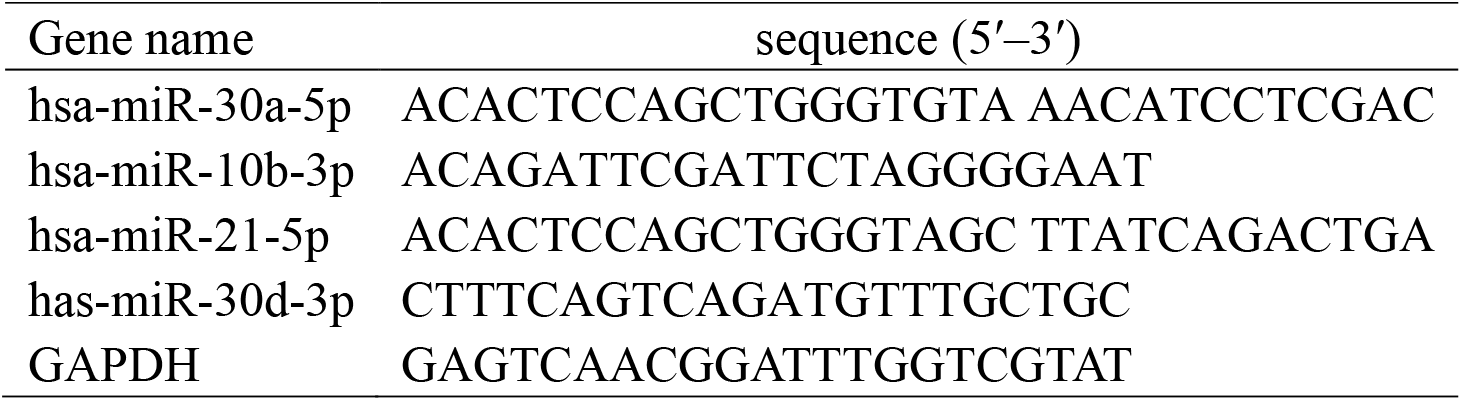
Primers used for RT-qPCR analysis

### Cisplatin-induced AKI model establishment

A cisplatin-induced acute kidney injury (AKI) model was established by using C57BL/6J female mice (8 weeks old, 20-25g). The mice were purchased from Zhejiang Vital River Laboratory Animal Technology Co.Ltd. (certificate number: 20200909Abzz0619000370). After intraperitoneal injection of pentobarbital (4 mg/kg) for anesthesia, 20 mg/kg of cisplatin was intraperitoneally injected into 8-week-old mice to establish AKI model[20]. Model group: 20 mice, 20mg/kg cisplatin were intraperitoneally administered according to the weight of the mice. Normal control group: 10 mice, same dose of normal saline were injected into the mice. Twenty-four hours later, blood was collected from the orbit. When the serum creatinine was significantly increased and 24-hour urine output significantly reduced, the model of acute kidney injury was successfully established. The mice in the model group were randomly assigned into AKI group (cisplatin group) and AKI+ exosomes group with 10 mice in each group. On the first day after the model, 100ug urinary exosomes from preterm infant (EVs) dissolved in 200uL PBS solution or the same volume of PBS were injected into the mice via tail vein. The mice were killed 72 hours after EV administration. Blood was collected to evaluate parameters of kidney function. The kidneys were preserved for histopathology and molecular analysis. The scoring of renal tubular damage was conducted according to Paller’s scoring standard[21].

This study was approved by the Animal Ethics Committee of the First Affiliated Hospital of Jinan University. All the experiments were conducted in strict accordance with the policies of the First Affiliated Hospital of Jinan University. Animals were bred in SPF (Specific pathogen Free, SPF) animal house without pathogenic microorganisms. The animal house was facilitated with appropriate temperature and humidity. The animals had free access to water and food.

### Cisplatin-induced HK2 cell model establishment

Human proximal renal tubular epithelial cells (HK2) were purchased from ATCC (ATCC®CRL-2190) and cultured in DMEM (Gibco) supplemented with 10% FBS (Moregate), 500 U/ml penicillin and 500 μg/ml streptomycin (Gibco). Cisplatin (1mg/ml, aladdin, China). The cells were placed in the 37°C incubator with 5% CO2. When the confluence reached 80%, the cells were inoculated in a 6-well culture plate for cisplatin stimulation and exosome intervention experiments . Time selection: According to the actual situation of this study, the cells were first stimulated by different concentrations of cisplatin (10mM, 20mM) to detect the apoptosis of HK2 cells[22]. Then, 25ug exosomes were used to treat the cells to detect the effects of exosomes on the expression of apoptosis-related protein in HK2 cell.

### Deoxynucleotide terminal transferase dUTP-mediated nick end labeling (TUNEL) detection

Kidney paraffin sections were stained by using the in-situ apoptosis detection kit (C1086, Beyotime) according to the instructions of the kit. Each section was fully deparaffinized, treated with proteinase K in 37°C for 30 minutes, and treated with 3% hydrogen peroxide for 5 minutes. The slides were incubated with TUNEL reaction mixture in 37°C for 1 hour. Then, the buffer was added and incubated with the slides in 37°C for 30 minutes to terminate the reaction. By using the microscope, three randomly selected TUNEL-positive cells were counted for quantitative analysis.

### Cell transfection and dual luciferase reporter gene detection system

HEK293T cells were inoculated in a 24-well plate. When the confluence reached 70-80%, miRNA and 3’UTR vectors were co-transfected into the cells following the instructions of LipofectaminRNAiMax. The culture medium was changed 6 hours after transfection. The fluorescein intensity was measured 48 hours after transfection. The reporter gene was labeled with Renilla luciferase, and firefly luciferase was used as the internal control. The ratio of Renilla luciferase activity to firefly luciferase activity in each well was calculated. Three replicate holes were set for each group.

### Statistical analysis

GraphPad Prism V8.01 was used to analyze the data, which were expressed as mean ± SE. T test was used to compare data between normally distributed groups, and one-way ANOVA were performed for comparisons of data with more than two groups.The analysis of variance and Dunnett were used to compare results between multiple groups. Image j software was used to analyze the results of Western blot. P value<0.05 was considered as statistically significant.

## Results

### Isolation of exosomes from urine samples of premature infants

Differential ultracentrifugation was used to separate the urine of premature infants (n = 20) to obtain exosomes. Under the transmission electron microscope, the exosomes derived from urine of premature infants were round or oval vesicle-like, and the size was nanoscale (Figure 1a). The average diameter of uEVs from premature infants detected by Nanoparticle Tracking Analysis (NTA) is 124±4 6.3nm.The expression of exosome markers, CD63, CD9, and CD81, were detected by Western Blot. (Figure 1b-c). In addition, as confirmed by flow cytometry analysis, exosomes expressed aquaporin (Aquaporin, AQP1), the surface marker of renal tubular cells, as well as exosome specific protein (CD9 and CD63) (Figure 1d).

**Figure1.**
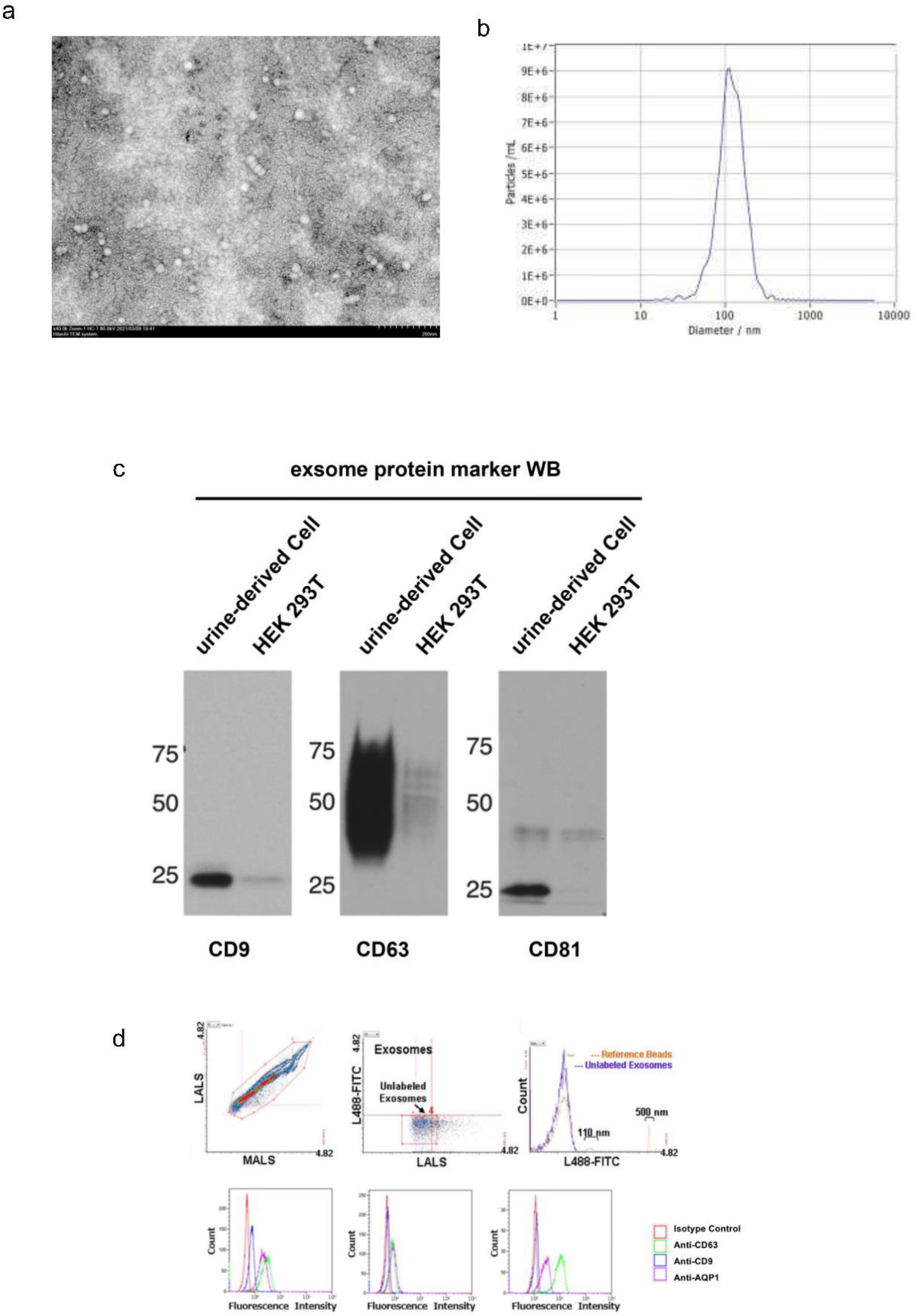
uEV Characterization: a:Representative micrographs of transmission electron microscopy obtained from uEVs (scale bars: 200 nm).b:Size analysis of uExo (mean diameter = 124.0 ± 46.3 nm). c: Representative western blot showing exosomal markers CD9,CD63,CD81. HEK293T cells was analyzed as positive control. d:Representative cytofluorimetric analyses of uEVs showing the positive expression of renal and exosomal markers AQP1, CD9, CD63.

### The exosomes derived from urine of premature infants reduced cisplatin-induced renal tubular damage and cell apoptosis in AKI models, and inhibited local inflammation

One day after AKI induction, as the peak value of the injury appeared, AKI mice were randomly assigned into cisplatin group and cisplatin + exosomes group (10 mice in each group). Two groups of mice received the following treatments respectively. In AKI+exosomes group, 200ul PBS solution containing exosomes (about 100ug) derived from urine of premature infants was injected into the mice via tail vein. The mice in cisplatin group was injected with the same volume of normal saline. The histological examination of kidney sections in the cisplatin group presented obvious lumen distortion and expansion, formation of urinary cylinder, and necrosis of renal tubular epithelial cells (Figure 2b). In the mice treated with uEVs from premature infants, the pathological changes were significantly ameliorated, the urinary cylinder and the number of necrotic renal tubular epithelial cells were significantly decreased, and the tubular damage score was reduced, which confirmed uEVs had protective effects (Figure 2c,d). Creatinine level was significantly decreased in the mice injected with uEVs from premature infants compared with the model group on the 4th day (72 hours after treatment) (P<0.01, but still higher than the normal control group, Figure 2e). Compared with the control group, the survival rate of mice in the exosome treatment group was significantly higher than that in the control group (Figure 2f). These results indicated that exosomes derived from urine of premature infants could attenuate acute kidney injury and protect renal function in AKI mice model.

**Figure 2.**
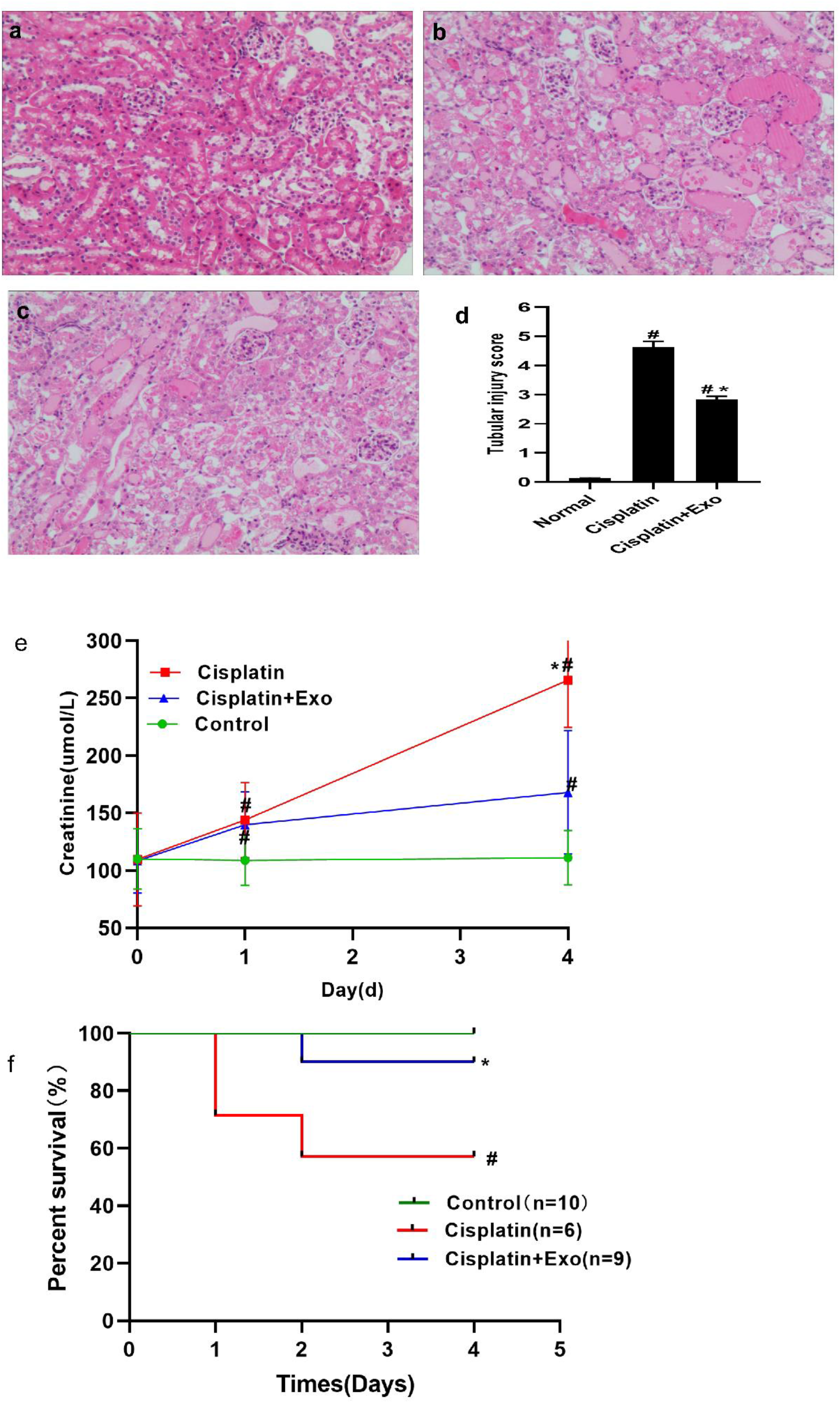
uEVs Improve Renal Recovery in an AKI Mouse Model. a-c:Representative micrographs of histological analysis (H&E staining) of renal tissue from healthy and AKI mice treated with saline uEVs at day 3 after damage. Original magnification 200.d:Quantification of tubular hyaline casts and tubular necrosis at day 3 after damage. Data are expressed as the mean ± SEM. Ten fields per section were analyzed.ANOVA test was performed: *p < 0.05 or #p <0.01 versus normal .e:Alterations in renal function were assessed by evaluating creatinine levels at day 3 after damage. Mice treated with uEVs isolated by ultracentrifuge (uEV), showed significantly reduced levels of creatinine compared with untreated AKI mice . Data are expressed as the mean ± SEM of eight mice per group. ANOVA test was performed: *p < 0.05 or _#_p < 0.001 versus vehicle.

We further analyzed the impact of uEVs on renal tubular injury. In particular, the expression level of caspase-3 was increased and that of Bcl2 decreased in AKI mice. These proteins were the markers for renal tubular epithelial cell injury. In contrast, in the mice treated with uEVs, the markers for injury and apoptosis, caspase-3 and Bcl-2, was significantly down-regulated and up-regulated, respectively (Figure 3a-d). In addition, a significant increase in pro-inflammatory cytokines, such as monocyte chemotactic factor (MCP1), was observed in the kidney tissue from AKI mice. Treatment with uEVs could inhibit the expression of the inflammatory marker, MCP1 (Figure 3e,f). By using TUNEL staining (Figure 4), the treatment with exosomes derived from urine of premature infants also significantly reduced the apoptosis of cisplatin-induced renal tubular epithelial cell on the 4^th^ day in the AKI model. Western blot showed that the expression of anti-apoptotic protein, Bcl2, was increased in mice treated with exosomes derived from urine of premature infants, while the expression of apoptotic protein, caspase-3, and inflammatory molecule MCP1 was decreased (Figure 5). In summary, these results indicated that treatment with urinary exosomes from premature infants ameliorated the apoptosis and inflammation, and promoted the repair process of renal tubular epithelial cell.

**Figure 3-4.**
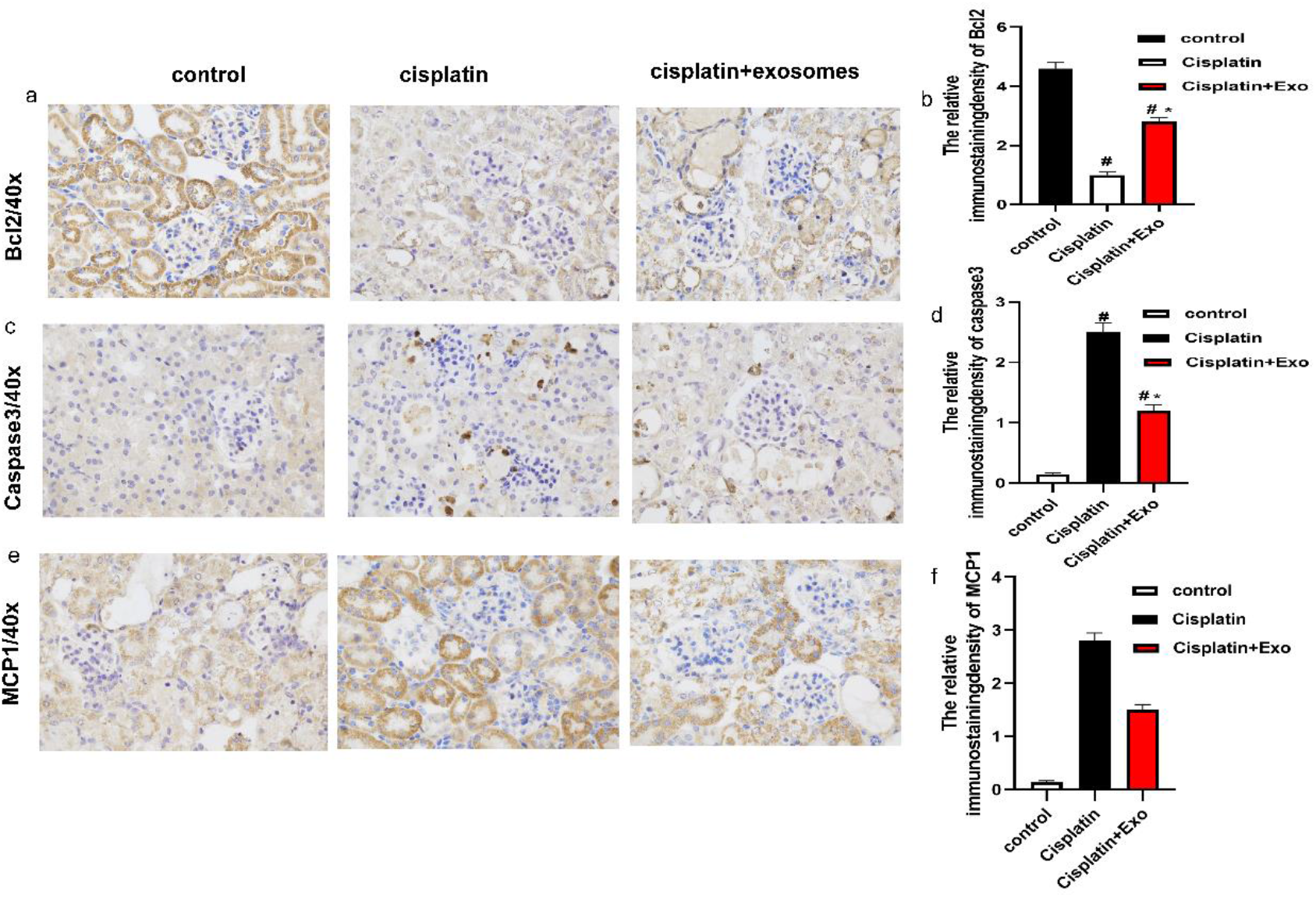
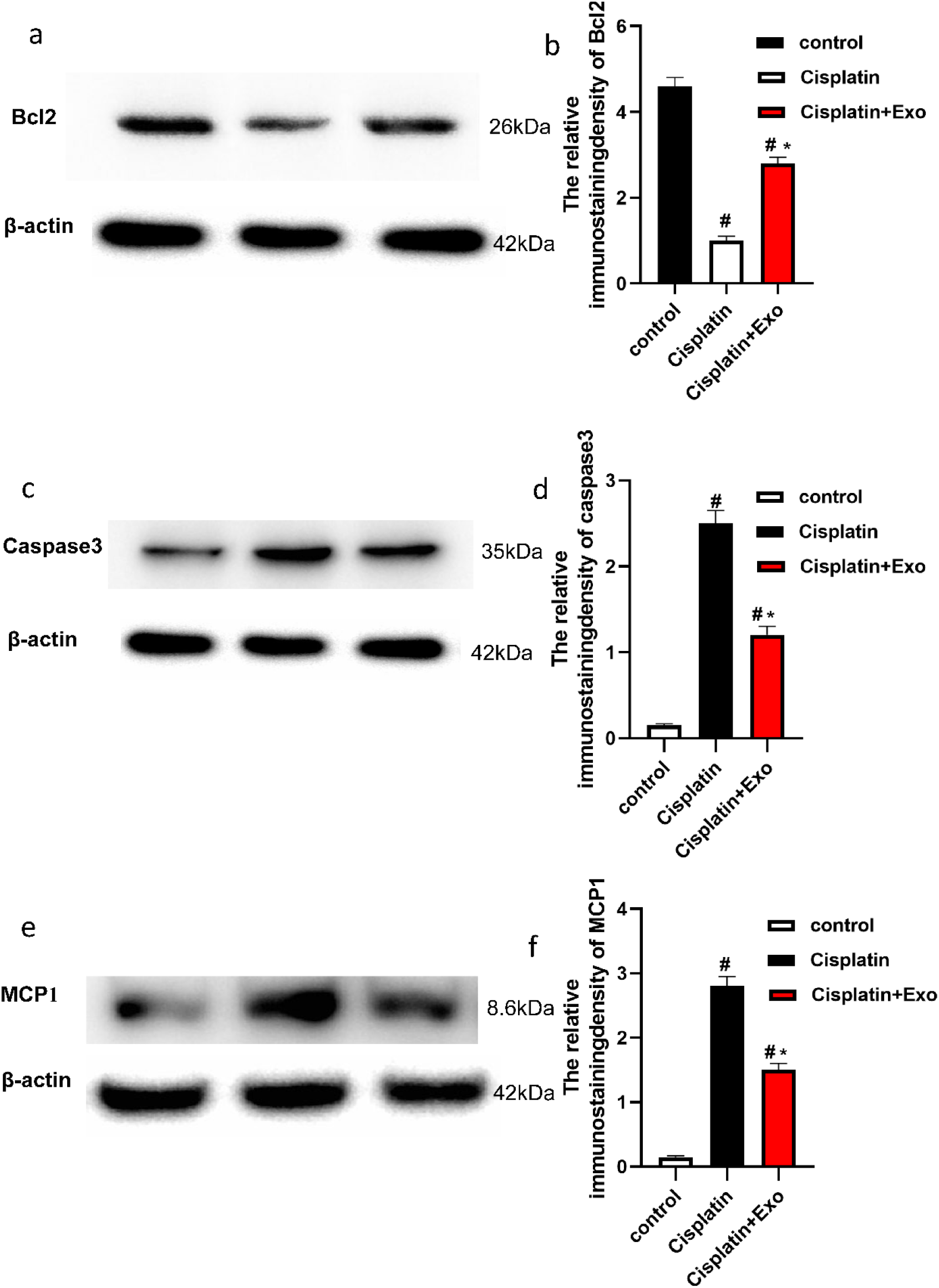
Exo could attenuate the expression of apoptosis-related protein. The relative protein levels of Bcl-2/ β-actin, caspase-3/β-actin MCP1 / β-actin were detected by Immunohistochemistry and Western blotting. _#_P < 0.05, versus normal; *P < 0.05, versus cisplatin.

**Figure 5.**
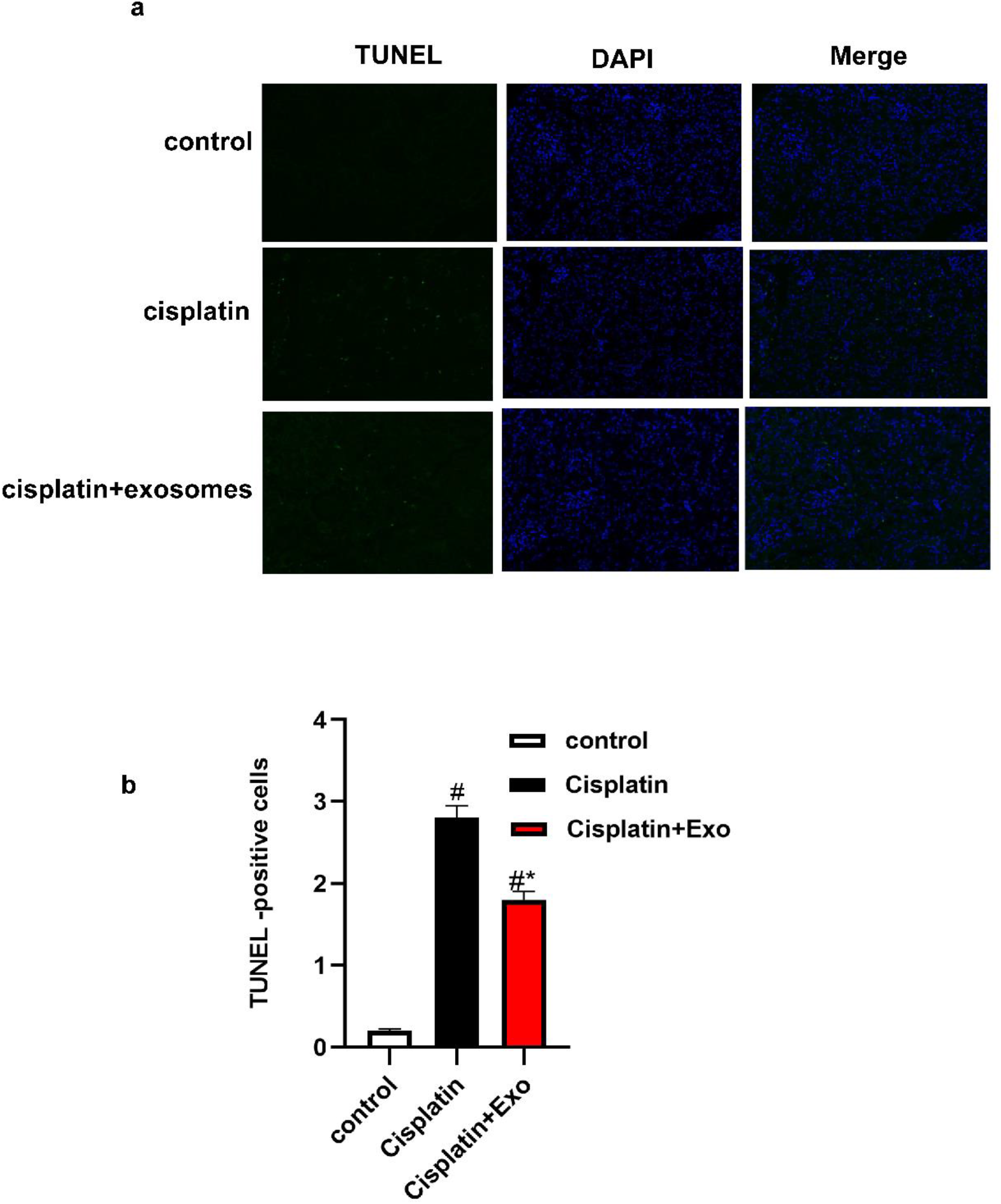
Exo could attenuate the TUNEL-positive cells. a:Tissue apoptosis was examined with TUNEL-positive staining.b: semi-quantitative analysis for apoptotic examination was scored.

### The exosomes derived from urine of premature infants inhibited HK2 cell apoptosis induced by cisplatin in vitro

The HK-2 cell line was treated with 10mmol/L and 20mmol/L cisplatin and 25ug/ml exosomes for 24h. Then, the apoptosis rate was detected by using Annexin V-FITC/PI double-labeling method. The apoptosis rate in the cisplatin group was significantly different from that of the control group (P<0.01). Compared with the cisplatin group, the apoptosis rate in the cisplatin+exosomes group was reduced from 49.5±4.4 (%) to 32.6±3.6 (%) with a significant difference (P<0.05) (Figure 6a-e). The experiment was repeated 3 times and the average value was calculated. Western blot analysis showed that treatment with urinary exosomes from premature infant reduced the expression of caspase-3 and Bax in HK2 cells induced by cisplatin, and increased the expression of Bcl2 (Figure 6-f). These results indicated that the urinary exosomes from premature infants had a protective effect on HK2 cell injury caused by cisplatin.

**Figure 6.**
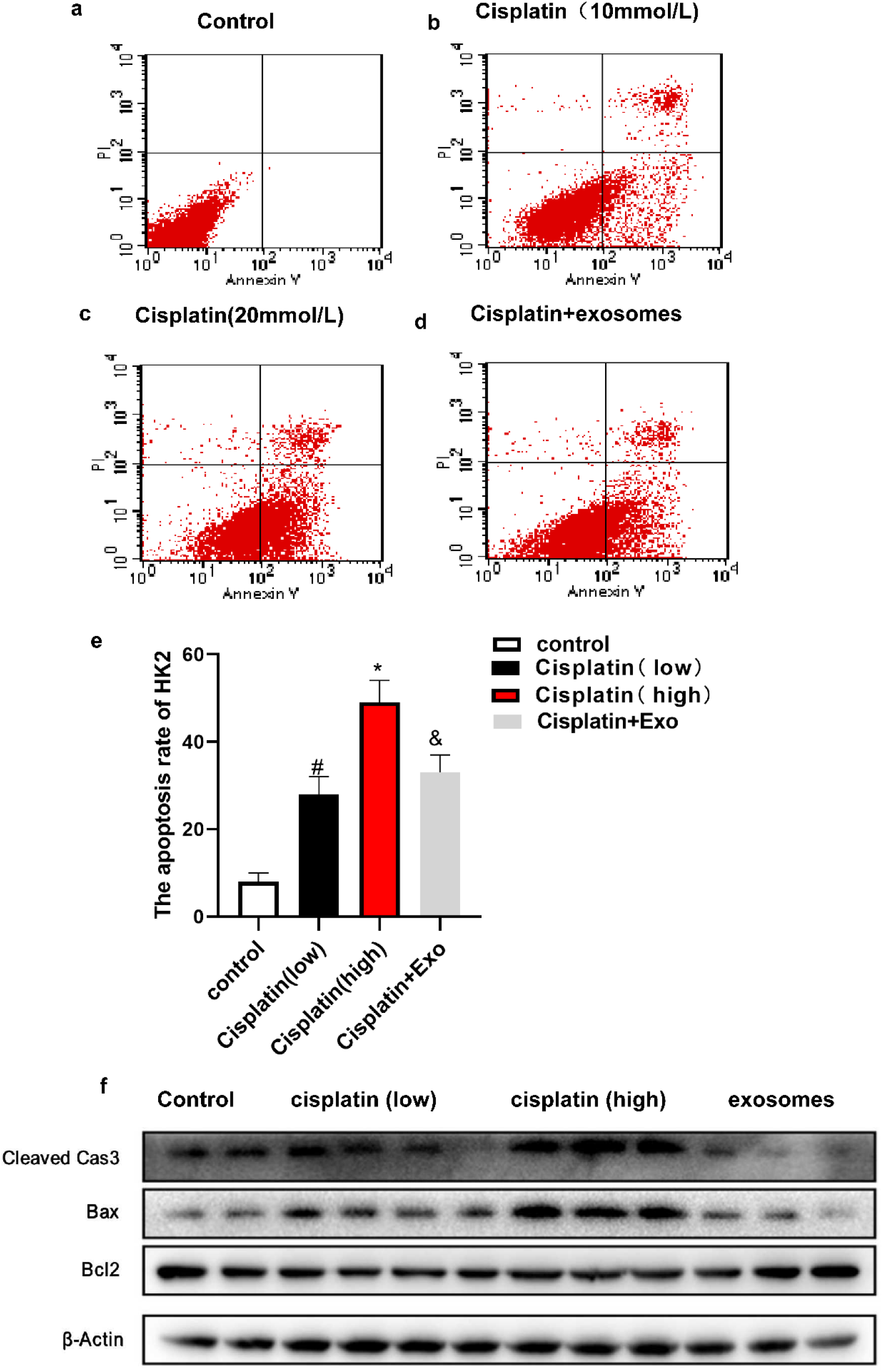
Exo could reduce apoptosis of human renal tubular epithelial cells induced by cisplatin. a-d) Representative FCM photograph of PI and AnnexinV-FITC double stained HK2 apoptosis. e) The bar chart of PI and AnnexinV-FITC double stained HK2 apoptosis. *P < 0.05, versus normal group; ^&^P < 0.05, versus treated with high cisplatin group.f) The relative protein levels of activated caspase-3,bax,and bcl2 were detected by Western blotting.

### The sequencing of miRNAs in the exosomes derived from urine of premature infants and the potential targets of miR-30a-5p

Previous research reported that exosome miRNAs were involved in a variety of physiological and pathological processes. In order to further investigate the mechanisms of the protective effect of urinary exosomes from premature infants, miRNAs in urinary exosomes from premature infants were sequenced. The results showed that 13 miRNAs were most enriched in urinary exosomes of premature infants (Figure 7a). We then validated the expression levels of the first four enriched miRNAs by qRT-PCR. It was determined that the expression level of miR-30a-5p was the highest among the miRNAs in urinary exosomes from premature infants(Figure 7b).. In order to further study the function of miR-30a-5p, bioinformatics tools Targetscan, and microT were used to predict the target genes of miR-30a-5p. There were 257 target genes were found to be the potential target genes of miR-30a-5p. By using GO and KEGG methods analyzing the target genes and the acute kidney injury model induced by cisplatin, the target gene MAPK8 was screened out, which was related to apoptosis or necrosis. By using TargetScan analysis software, it was found that miR-30a-5p had a binding site with the 3’UTR region of MAPK8 in mouse, which was located at 59-66nt, a conservative site of MAPK8 3’UTR (Figure 7c). After constructing the vectors containing wild-type and mutant MAPK8 3’UTR and miR-30a-5p, the vectors were transfected into HEK293T cells to detect Renilla and firefly luciferase activities. The results showed that compared with the group of co-transfection with the plasmids containing 3’UTR-NC and miR-30a-5p, the ratio of Renilla luciferase /firefly luciferase activity was significantly decreased in the group of co-transfection with the plasmids containing MAPK8 3’UTR and miR-30a-5p (p<0.01). In the group of co-transfection with the plasmids containing mutant MAPK8 3’UTR and miR-30a-5p, the ratio was basically the same as that of the control group (p>0.05, Figure 7d). This confirmed that miR-30a-5p could bind with the 3’UTR region of MAPK8. In order to further prove whether miR-30a-5p could act on MAPK8, Western Blot was used to detect the change of MAPK8 expression in HK2 cells after stimulated by miR-30a-5p mimic. In Figure 8d, the expression of MAPK8 was decreased after the cells were stimulated by 10nM miR-30a-5p mimic (p<0.05). These results indicated that miR-30a-5p in urinary exosomes from premature infants might bind with the 3’UTR region of MAPK8 mRNA, thereby inhibiting the expression of MAPK8 via post-translational modulation.

**Figure 7.**
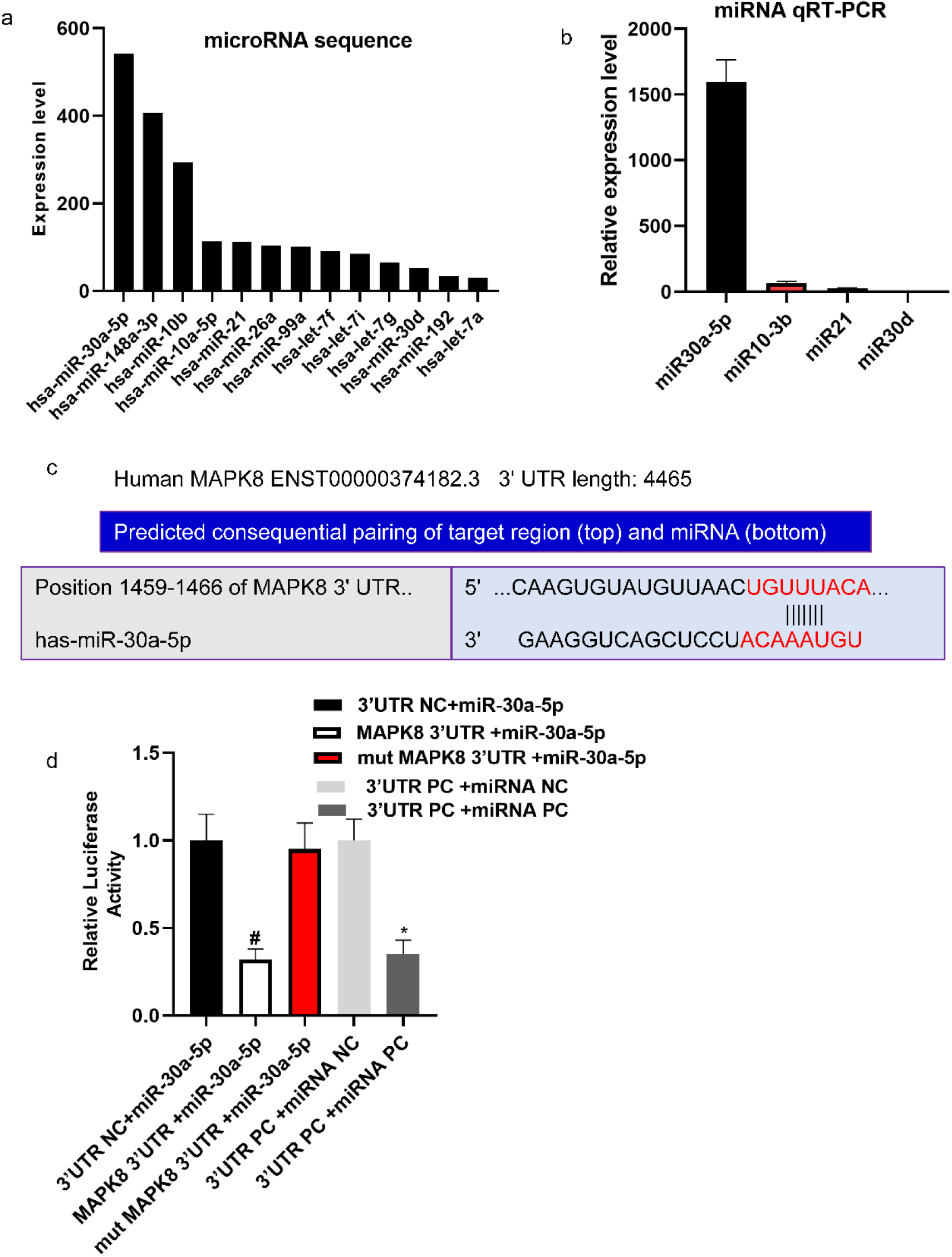
miRNA sequencing of Exo contents and potential targets of miR-30a-5p. a:the top 13 most-enriched miRNAs in Exo. b: qRT-PCR analysis of the expression levels of the top 4 most-enriched miRNAs in Exo. **c**:Sequence alignments of miR-30a-5p and its candidate target sites in the 3’UTR of MAPK8. **d**: Luciferase reporter assay of miR-30a-5p mimic-treated HEK293T cells, which suppresses MAPK8-wildtype 3’UTR. Data represent the mean ± SEM. *P<0.05, ^#^P<0.05.

**Figure 8.**
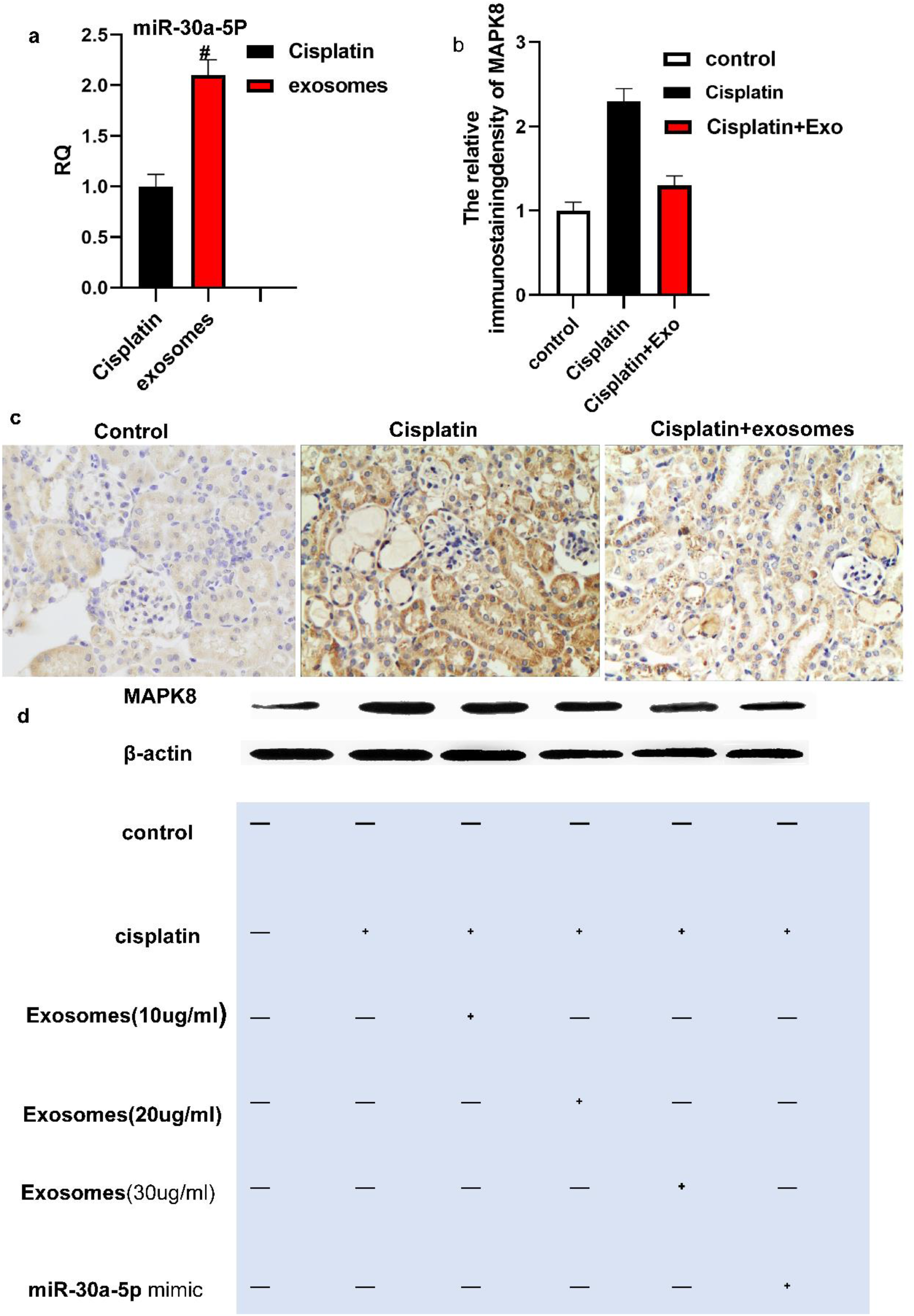
uExo upregulate miR-30a-5p expression, which targets the MAPK8 in vivo and in vitro. a: qRT-PCR analysis of the relative expression levels of miR-30a-5p (P=0.01) b:.semi-quantitative analysis for MAPK8 expression was scored. c:immunohistochemistry analysis of MAPK8 expression in the control group (n=10) ,AKI group (n=6) and AKI treated with uExo group (n=9) on day 3 after AKI. d: uExo or miR-30a-5p mimic reduces MAPK8 expression in cisplatin induced injury of HK2 cells.

### The exosomes derived from urine of premature infants enhanced miR-30a-5p expression and inhibited MAPK8

In order to further confirm that miR-30a-5p was essential to the protective effects of urinary exosomes from premature infants on AKI, we first determined the expression level of miR-30a-5p in AKI kidney tissues treated by exosomes. qRT-PCR analysis demonstrated that the expression level of miR-30a-5p was increased in the kidney tissue from AKI mice treated by exosomes (Figure 8a). The results of immunohistochemistry further proved that the expression of MAPK8 was down-regulated in the kidney tissue from AKI mice treated by exosomes (Figure 8b-c). In vitro, urinary exosomes from premature infants and miR-30a-5p mimic inhibited the expression of MAPK8 in HK2 cells induced by cisplatin (Figure 8d).

## Discussion

In this study, we evaluated the role of the urinary exosomes from premature infants in renal regeneration and repair in vivo and in vitro. In the experimental mouse AKI model induced by cisplatin, the treatment with urinary exosomes from premature infants significantly reduced mortality, ameliorated renal function and histological abnormalities, and promoted the recovery of acute kidney injury. We further demonstrated that urinary exosomes from premature infants had the effects of anti-apoptosis and inhibiting inflammation activation, which were similar to the protective effects of mesenchymal stem cell-derived exosomes on AKI observed in past research[8]. In particular, our study revealed that urinary exosomes from preterm infants could down-regulate the expression of MAPK8 via delivering miR-30a-5p to inhibit renal tubular epithelial cell apoptosis and promote the repair of renal tubular epithelial function. In addition, treatment with urinary exosomes from premature infants significantly inhibited the activation of pro-inflammatory cytokines during acute kidney injury.

In the past, AKI was considered to be a self-limiting disease. Recent studies supported that the acute change in renal function was associated with long-term prognosis, including progression to chronic kidney disease and cardiovascular disease, persistent dysfunction, and even death[1]. The pathophysiological response to AKI may determine whether kidney function can be restored or progress to chronic kidney disease (CKD).The pathophysiological mechanisms involved in the injuries caused by various etiologies are different. The common mechanisms include abnormal hemodynamics, renal tubular epithelial cell death, oxidative stress injury, renal tissue inflammation, and immune response[23].Among them, renal tubular epithelial cell damage is the initial and key part that causes AKI. The damage of renal tubules may be directly caused by various damaging factors, and may also be secondary to glomerular or vascular disease[24, 25].Usually, renal tubular epithelial cells (TEC) have a strong self-renewal ability. When the source of damage is removed, TEC can quickly repair. However, in recent years, accumulating studies have shown that renal tubular cell not only acted as a passive victim, but may also be activated by TEC to synthesize and release various biologically active factors, recruit inflammatory cells to accumulate in the tubulointerstitium, amplify the inflammatory cascade, and lead to TEC apoptosis, and thereby resulting in death, pyrolysis, necrosis, and increased extracellular matrix production[26]. This vicious circle of necrotizing inflammation eventually leads to renal dysfunction and fibrosis [27].In addition, unrepaired TEC can cause cell cycle block (G2/M phase block). Blocked TECs secrete a large number of factors such as TGF-β and CTGF, and promote the formation of renal fibrosis [24, 28].There is a lack of effective clinical measures to reverse this process. The potential curative effect of extracellular vesicles derived from mesenchymal stem cell, especially exosomes, on kidney disease has been widely reported. It is considered to be a new non-invasive therapeutic strategy for kidney regeneration, which may act through a variety of regeneration processes, including inducing the survival and proliferation of renal tubular cell, inhibiting inflammation and fibrosis [18]. and so on. However, due to the invasive features of the collection process and limited availability of stem cells and exosomes, wide application of this method is limited. In addition, studies have shown that MSC EV could not express nephroprotective protein Klotho. Renal-derived EVs isolated from normal urine contained Klotho molecules, which could protect against acute kidney injury caused by glycerol injection. The mechanism was that uEV stimulated the proliferation of renal tubular cells, reduced the expression of inflammation and injury markers, and compensated the loss of endogenous Klotho [17]. EVs secreted by cultured renal tubular cells can restore ischemia-reperfusion kidney injury. Urinary exosomes can preferentially locate in the kidney injury area to transfer specific miRNA. The analysis of the targets of uEV-miRNA interaction revealed that the activation of growth factor pathways such as IGF-1 further promoted cell regeneration[17]. Therefore, the mechanism of urinary exosomes in repairing kidney damage might not be same as that of the mesenchymal stem cell exosomes, and urinary exosomes are more kidney-targeting. In this study, we aimed to explore the effect of urinary exosomes from premature infants on cisplatin-induced acute kidney injury and its possible mechanism.

Previous studies have proved the effectiveness of exosomes in the stem cell derived from adult urine in the repair of acute kidney injury. It was reported that intravenous injection of urinary stem cell exosomes (USCs-exosomes) in healthy adults could reduce urinary microalbumin excretion rate in diabetic rats, prevent podocytes and renal tubular epithelial cells from apoptosis, inhibit caspase-3 overexpression, and increase the proliferation of glomerular endothelial cells [24]. In addition, USCs-exosomes can reduce cell apoptosis induced by high glucose in vitro [27]. The mechanism may be that USCs exosomes contain potential factors such as growth factors, transforming growth factor (TGF-β1), angiopoietin, and bone morphogenetic protein-7, which may be related to angiogenesis and cell survival [28]. Recent studies have found that kidney-derived EVs isolated from normal human urine contain Klotho molecule, which could protect renal function. Kidney-derived EVs could protect mice from acute kidney injury caused by glycerol injection. The mechanism was that uEV stimulated the proliferation of renal tubular cells and reduced the expression of the markers for inflammation and injury, and compensated the loss of endogenous Klotho [17]. In addition, a recent study discovered that stem cell exosomes derived from fresh adult urine contained abundant miR-146a-5p, which could inhibit the activation of NF-κB signaling pathway and the infiltration of inflammatory cells by targeting the 3’UTR of IRAK1 and served a protective effect on blood reperfusion injury [18]. In summary, healthy adult urine-derived exosomes may be a new therapeutic strategy for renal-derived regeneration. The transfer of the biological components (protein, RNA and DNA) of urine-derived EV to target cells seems to be the main mechanism in the function of EVs[13]. However, there is a lack of studies on the effects of urinary exosomes from premature infants on renal injury. In our experiments, the protective effect of urinary exosomes from premature infants on cisplatin-induced AKI was consistent with the previous studies. Our results suggested that urinary exosomes from premature infants could improve the significant decrease of serum indexes of renal function such as creatinine, reduce mortality, significantly ameliorate renal histological damage caused by cisplatin, and decrease the score of renal tubular damage. Meanwhile, the results of immunohistochemistry presented that urinary exosomes of premature infants significantly reduced the expression of MCP-1, Caspase-3 and other inflammatory genes in acute kidney injury tissues, and increased the expression of anti-apoptotic gene, Bcl2. In consistent with these results, urinary exosomes from premature infants could also protect cultured HK2 cells from cisplatin injury in vitro. In brief, the treatment with urinary exosomes from premature infants accelerated the recovery of the kidney, ameliorated the abnormal function and histological damage, and inhibited the apoptosis of renal tubular cells.

MicroRNAs, a type of small non-coding RNAs with about 17-24 nt in length, can mediate post-transcriptional gene silencing by binding to the 3′-untranslated region (UTR) or open reading frame (ORF) region of the target mRNA. miRNAs play a key regulatory role in various pathophysiological activities, including cell pro liferation, differentiation, migration, disease occurrence and progression[15]. It is estimated that the human genome encodes more than 1,000 miRNAs, targeting about 60% of human protein - coding genes[29]. As the transportability of vesicles is widely known, the role of miRNAs in exosomes has attracted more and more attention. Through mass spectrometry and array analysis, more than 41000 unique gene products and more than 2800 miRNAs have been identified in EVS[12, 30]. Therefore, we performed miRNA sequencing on urinary exosomes from premature infants. An important finding was that miR-30, miR-10, and let-7 family members presented the most abundant expression. This was consistent with previous research results[19]. It was determined that miR-30a-5p was the most abundant miRNA. The role of miR-30a-5p in kidney disease has been reported. It was reported that miR-30a-5p inhibited renal hypoxia/reoxygenation (H/R)-mediated apoptosis of HK2 cells[31]. Accumulating studies have found that in the glomeruli of patients with glomerular diseases including DN, FSGS or MPGN, the expression of miR-30a-5p was down-regulated, especially in podocytes [32–34].The level of urinary miR-30a-5p before treatment could not only distinguish the CR group from the non-CR group, but also predicted the response to hormone therapy[33]. Shihana F et al. found that among the types of AKI caused by nephrotoxic drugs, there were 7 kinds of microRNAs that could distinguish patients with severe AKI from those without AKI, and there is a change of more than 17 times in four of them (miR-30a-3p, miR-30a-5p, miR-92a, and miR-204)[35]. In this experiment, we detected the change of miRNA expression in the kidney tissue of AKI mice on the 4th day after exosome treatment. It was found that the expression of miRNA-30a-5p in the kidney tissue was increased significantly after exosome treatment, suggesting that miRNA-30a-5p might play an important role in the repair of renal injury, which could be a new target for the treatment of renal injury and worthy of further study.

We further studied the downstream mechanism of miR-30a-5p in the urinary exosomes from preterm infants. The results showed that it could down-regulate the expression of MAPK8. MAPK8, also named as Jun amino-terminal kinase (JNK), is a stress-activated protein kinase (SAPK). It is a member of the MAPK family and can be stimulated by a series of extracellular stimuli including inflammatory cytokines and physiological stress, such as growth factors and GPCR agonists. It can regulate cell proliferation, differentiation and survival. Previous studies confirmed that cisplatin could activate p38, ERK and MAPK8 signaling pathways in renal tubular epithelial cells [20]. Continuous activation of the MAPK pathway directly promoted the death of renal tubular epithelial cells, and mediated the up-regulation of the pro-apoptotic gene Bax in mitochondrial-dependent apoptosis signaling pathways, which in turn led to the release of cytochrome c and activation of caspase-3 [36]. In addition, MAPK8-mediated pathway activation not only participated in G2/M block and apoptosis of renal tubular epithelial cell ,but also up-regulated the expression of TGF-β1 and CTGF in renal tubular cells with G2/M block, leading to abnormalities in renal tubular epithelial cells repair, production of collagen, and promoted the development of fibrosis [13]. This study not only confirmed that MAPK8 was involved in the damage of renal tubular epithelial cells caused by cisplatin, but more importantly, we revealed that urinary exosomes down-regulated the expression of MAPK8 through miR-30a-5p targeting, inhibited MAPK signaling pathway, and promoted kidney repair. Studies have found that miR-30a attenuated acute kidney injury by up-regulating the expression of Klotho protein[37]. In addition, miR-30a-5p also directly targeted CTGF to inhibit the formation of myocardial extracellular matrix [38]. Combined with current results, targeting miR-30a-5p and its target genes may be a potential therapeutic way for the treatment of kidney disease.

There were some limitations in our research. First of all, due to the limitations of our experimental conditions, it was impossible to specifically label, track and detect the release of urinary exosomes from preterm infants in vivo and their enrichment in the kidneys. This design could more directly prove that the urine from preterm infants protect renal function via exosomes. Secondly, there are a large number of miRNAs in the urinary exosomes from premature infants. These miRNAs may play a potential regulatory role in the development of the kidney and repair of damage, which is supported by bioinformatics research. Dynamic evaluation of miRNAs released by exosomes and changes in related target pathways can benefit the development of non-invasively detecting AKI and find key targets in disease progression.

## Conclusions

In summary, we have proved that urinary exosomes from premature infants could protect cisplatin-induced acute kidney injury in mice, and had protective effects such as anti-apoptosis and anti-inflammatory functions. Moreover, by using the cisplatin-induced HK2 cell in-vitro model, it was determined that miR-30a-5p in urinary exosomes from premature infants mediated cytoprotective effects via down-regulating MAPK8. These findings provided a theoretical basis for the use of urine-derived exosomes from premature infants to treat AKI.

## Abbreviations

AKI: acute kidney injury
HK2: human kidney cortex/proximal tubule cells
USC-Exo: exosomes derived from human urine-derived stem cells
CKD: Chronic kidney disease
qRT-PCR: quantitative reverse-transcriptase polymerase chain reaction
TUNEL: Terminal deoxynucleoitidyl transferase-mediated nick end labeling
sCr: serum creatinine
MAPK8: mitogen-activated protein kinases-8
ESRD: end stage renal disease
EVs: extracellular vesicles
MSC: mesenchymal stem cells
hUSCs: human urine-derived stem cells
PBS: phosphate buffer saline
TEM: transmission electron microscope
SPF: Specific pathogen Free
NTA: Nanoparticle Tracking Analysis
TEC: tubular epithelial cells

## Study approval

The institutional review board in the First Affiliated Hospital of Jinan University approved this study. The experimental protocol were approved by the administrators of the management of scientific research office of First Affiliated Hospital of Jinan University in Jinan University involving human subjects,informed consent was obtained from all participants.

## Consent to publish

All authors have read and approve the final version of the manuscript. All authors agree with submission and consent to publish.

## Author contributions statement

MMM and QL design, collection and assembly of data, data analysis and interpretation, manuscript writing. LJF, HWW, and YPL animal model and collection and assembly of data, generated the data for the manuscript. YM, SC, BH FNL, BH,and BZG, experiment instruction, technique support, and edited the manuscript. HH,L,SLH,and WXL collection and assembly of data. LH Y and YFH design, financial support, result interpretation, edited the manuscript, final approval of manuscript.

## Acknowledgements

This study was supported by the Guangdong Academician Workstation, Industry-University-Research Cooperation (Grant no. 2013B090400004).

